# Salicylic acid mediated immune response of *Citrus Sinensis* to varying frequencies of herbivory and pathogen inoculation

**DOI:** 10.1101/2020.08.04.235911

**Authors:** Freddy Ibanez, Joon Hyuk Suh, Yu Wang, Monique Rivera, Mamoudou Setamou, Lukasz L. Stelinski

## Abstract

Plant immunity against pathogens and pests is comprised of complex mechanisms orchestrated by signaling pathways regulated by plant hormones [Salicylic acid (SA) and Jasmonic acid (JA)]. Investigations of plant immune response to phytopathogens and phloem-feeders have revealed that SA plays a critical role in reprogramming of the activity and/or localization of transcriptional regulators via post-translational modifications. We explored the contributing effects of herbivory by a phytopathogen vector [Asian citrus psyllid, *Diaphorina citri*] and pathogen [*Candidatus* Liberibacter asiaticus (*C*Las)] infection on response of sweet orange [*Citrus sinensis* (L.) Osbeck] using manipulative treatments designed to mimic the types of infestations/infections that citrus growers experience when cultivating citrus in the face of Huanglongbing (HLB) disease. A one-time (7 d) inoculation access period with *C*Las-infected vectors caused SA-associated upregulation of *PR-1*, stimulating defense response after a long period of infection without herbivory (270 and 330 days). In contrast, while repeated (monthly) ‘pulses’ of 7 d psyllid feeding injury stimulated immunity in CLas-infected citrus by increasing [SA] in leaves initially (up to 120 d), long-term (270 and 330 days) repeated herbivory caused [SA] to decrease coincident with upregulation of genes associated with SA metabolism (*BMST* and *DMR6*). Similarly, transcriptional responses and metabolite (SA and its analytes) accumulation in citrus exposed to a continuously reproducing population of *D. citri* exhibited a transitory upregulation of genes associated with SA signaling at 120 days and a posterior downregulation after long-term psyllid (adults and nymphs) feeding (270 and 330 days). Herbivory played an important role in regulation of SA accumulation in mature leaves of *C. sinensis*, whether or not those trees were coincidentally infected with *C*Las. Our results indicate that prevention of feeding injury inflicted by *D. citri* from the tritrophic interaction may allow citrus plants to better cope with the consequences of *C*Las infection, highlighting the importance of vector suppression as a component of managing this cosmopolitan disease.

**Author Summary:** We explored tritrophic interactions among an insect vector (*Diaphorina citri*) – phytopathogen (*Candidatus* Liberibacter asiaticus) – and cultivated fruit crop [sweet orange, *Citrus sinensis* (L) Osbeck]. Transcriptional and metabolic responses of plants were analyzed over an extended time-course of disease progression after various frequencies of herbivore feeding and durations of pathogen infection using manipulative treatments designed to mimic the types of infestations/infections that citrus growers experience when cultivating citrus in the presence of the devastating citrus disease, huanglongbing. We found that in the absence of coincident psyllid feeding damage, citrus trees could mount a defense response against the pathogen by activating the salicylic acid (SA) pathway. Repeated, monthly ‘pulses’ of herbivory led to pronounced stimulation of SA transcription that was coincident with diminished pathogen titers in plants. Although insect injury initially activated SA-dependent defense responses, continuous and/or long-term (≥ 270 d) herbivory shut down *PR-1*-dependent defense responses against the pathogen. Our results provide a mechanism explaining how vector suppression contributes to maintaining health of cultivated citrus in areas where huanglongbing is endemic. Our results also point to specific gene targets that may yield novel genotypes expressing tolerance against *C*Las after appropriate manipulations.

## Introduction

Huanglongbing (HLB), also known as citrus greening is a disease that limits production of commercially important *Citrus* spp. worldwide [1]. Three taxonomic species of pathogenic bacteria are associated with HLB: *Candidatus* Liberibacter asiaticus, *C.* Liberibacter africanus, and *C.* Liberibacter americanus. In Florida, only *C.* Liberibacter asiaticus (*C*Las) has been reported and is transmitted by the vector, Asian citrus psyllid, *Diaphorina citri* Kuwayama [1, 2]. Despite efforts to control the disease and/or vector, current strategies have had limited effect on curtailing disease spread resulting in lost commercial feasibility. Florida’s citrus industry has been devastated losing over $7 billion [3], putting the future of America’s citrus production capabilities at risk. Nevertheless, significant research effort targeting this problem is beginning to yield promising results, including identification of two varieties that exhibit levels of tolerance to *C*Las infection - ‘Bearss’ lemon, *Citrus limon* Burm. f., and ‘LB8-9’ Sugar Belle^®^ mandarin hybrid (“Clementine” mandarin × “Minneola” tangelo). Lower levels of phloem disruption and greater phloem regeneration have been documented in these two varieties following *C*Las infection as compared with HLB-susceptible sweet orange, *Citrus sinensis* (L.) Osbeck [4]. However, the molecular mechanisms involved in this tolerance are still not understood and the biological events involved in defense systems of *Citrus* varieties require further investigation.

Plant defense mechanisms are an adaptive response to combat injurious organisms. Two innate immunity strategies *in planta* have evolved to detect pathogens and their mechanisms have been extensively reviewed [5–7]. The first strategy (barrier) is known as microbe-associated molecular patterns (MAMPs) or pathogen-associated molecular patterns (PAMPs) [8–10] wherein pattern-recognition receptors localized in the host-cell membrane interact with conserved molecules of different groups of microbes, including: lipids, carbohydrates, proteins, and other small molecules. This recognition activates PAMP-triggered immunity (PTI) leading to an array of defense responses [5]. The second strategy, effector-triggered immunity (ETI), involves the secretion of pathogen effectors that interplay with host proteins commonly called [resistance (R) genes]. Successful pathogens that suppress host PTI responses, for instance the hypersensitive response, secrete specific effectors that modulate programmed cell death activated during PTI [11]. Most R genes encode nucleotide-binding site, leucine-rich, repeat (NB-LRR) proteins [12, 13]; for example, BRASSINOSTEROID INSENSITIVE 1-ASSOCIATED KINASE 1 (BAK1), which is targeted by bacterial effectors to interrupt the PTI response [14–16]. Recognition of bacterial molecules by PAMP or MAMP pattern-recognition receptors induces an oxidative burst produced by NADPH oxidase encoded by *rbohD* [17]. Also, such recognition activates the salicylic acid (SA)-dependent signaling pathway [18].

Plants not only must defend themselves against potential pathogens, but also encounter numerous herbivores during their lifetime that modulate specific biochemical and molecular responses, which vary depending on the type of feeding and/or oviposition occurring on the plant [19]. Investigations of plant-insect interactions using aphids and whiteflies as models have described induction of the SA-dependent immune response in plants as reviewed by Walling (20) and references therein. Similar investigations focusing on psyllids have demonstrated congruent immune responses in plants [21–23].

Plants synthetize SA using isochorismate synthase and phenylalanine ammonia-lyase enzymes and their respective pathways; these processes are reviewed in Chen, Zheng (24) and D’Maris Amick Dempsey, Vlot (25) and references therein. Recently, a non-enzymatic reaction for SA accumulation was described in *Arabidopsis thaliana*, wherein avrPphB Susceptible 3 (PBS3) in the cytosol catalyzes the formation of Isochorismate-9-glutamate (ISC-9-Glu), which later spontaneously decays into SA [26]. Afterward, SA is enzymatically modified by glycosylation, methylation, amino acid conjugation, and hydroxylation [25, 27]. The function(s) of SA metabolites have not been completely elucidated; but, putative roles were ascribed in defense regulation [28].

During SA-dependent signaling, proteins with major immune defense roles have been described; among them, the transcription cofactor, Nonexpresser of Pathogenesis Related genes 1 (NPR1), is a critical protein in plant defense during SA-dependent signaling. After nuclear translocation, NPR1 cofactor interacts with TGA transcription factors [29], and is proposed to function as a co-activator of gene expression for systemic acquired resistance (SAR) [30, 31]. Within the SA signaling pathway, two NPR1 paralogues, NPR3 and NPR4, were initially shown to interact with NPR1 and function as E3 ligases to degrade NPR1 in a [SA]-regulated manner to activate defense-gene expression [32, 33]. However, a recent study showed that NPR3/NPR4 interacts with TGAs to inhibit defense-related gene expression when [SA] is low. During pathogen infection, SA increases and binds to NPR3/NPR4, which releases the transcriptional repression of defense genes, allowing SA to bind NPR1, which in turn, activates the transcription of defense genes. We examined the mechanism(s) of immune response in *C. sinensis* via accumulation of SA and its metabolites after 7-, 14-, and 150-days of feeding by uninfected *D. citri*. Depending on the duration of herbivory, the vector differentially regulates transcription and accumulation of SA and its metabolites in mature leaves [23]. An increasing body of evidence suggests that immune responses against phloem-feeders and pathogens are tightly regulated by SA accumulation. We hypothesized that a threshold [SA] exists in *C. sinensis* that modulates the transcriptional regulation of genes involve in immune defenses and varies in accumulation in response to an interaction between stressors (pathogen and/or vector).

The overall goal of this study was to investigate modulation of SA-dependent plant defense responses in *C. sinensis* L. Osbeck (sweet orange). Three distinct frequencies of *D. citri* herbivory and *C*Las-inoculation access periods were imposed on *C. sinensis* as manipulated stressors. Our results describe the molecular and SA-dependent metabolic events that occur during HLB disease progression and after various intervals of insect feeding and following various frequencies of *C*Las inoculation. Specifically, we analyzed expression of a selected group of genes involved in: plant immune response, SA modification, and downstream SA signaling (perception). Concurrently, LC/MS analyses were conducted to quantify accumulation of SA and its metabolites. Based on our results, we propose working models to describe immune response in *Citrus* species following various intervals of *D. citri* infestation and frequencies of *C*Las inoculation.

## Results

### *C*Las detection in mature leaves of *C. sinensis*

The pattern of *C*Las-DNA titer in trees that were exposed to *C*Las-infected *D. citri* for one-time, 7-d inoculation access period (IAP) was similar to that found in trees that were exposed to a reproducing population of *C*Las-infected *D. citri* (presence of eggs, nymphs and adults) (Table 1). However, trees exposed to monthly pulses of 7 d IAPs using *C*Las-infected *D. citri* exhibited significantly lower *C*Las titer than the other two treatments for the duration of the experiment (12 months), with detectable levels of *C*Las DNA (0.010, 0.011 and 0.080 pg) at 240-, 330-, and 360-days, respectively. Based on these results, we chose time points when bacteria were detected in leaves (120-, 270-, and 360-days after first *C*Las-IAP) to analyze gene expression involved in SA-dependent defense response, as well as, accumulation of SA and SA-metabolites.

**Table 1.** Quantification of *C*Las genomic DNA in mature leaves of *C. sinensis* after initiation of *C*Las-IAP. The time-points chosen for gene expression and metabolite analyses were highlighted in bold, italicized, and underlined. Data represent mean ± standard deviation (SD) of six biological replicates per treatment. Different letters indicate statistical differences at *P* < 0.05. Abbreviation ‘N.D.’ denote No Detection of *C*Las DNA.

| pg of CLas genomic DNA per 10 g of *C. sinensis* DNA |  |  |  |
| --- | --- | --- | --- |
| Days after first CLas-IAP | One-time inoculation | Pulsed-inoculations | Continuous-inoculations |
| 30 | N.D. | N.D. | N.D. |
| 60 | N.D. | N.D. | N.D. |
| 90 | N.D. | N.D. | N.D. |
| **<u>120</u>** | 0.715 ± 0.172<sup>a</sup> | N.D. | 0.308 ± 0.130<sup>a</sup> |
| 150 | N.D. | N.D. | N.D. |
| 180 | N.D. | N.D. | N.D. |
| 210 | N.D. | N.D. | N.D. |
| 240 | N.D. | 0.010 ± 0.009<sup>b</sup> | N.D. |
| **<u>270</u>** | 0.680 ± 0.101<sup>a</sup> | N.D. | 0.721 ± 0.270<sup>a</sup> |
| 300 | N.D. | N.D. | 0.041 ± 0.021<sup>a</sup> |
| 330 | 0.662 ± 0.271<sup>a</sup> | 0.011 ± 0.010<sup>b</sup> | 0.818 ± 0.081<sup>a</sup> |
| **<u>360</u>** | 2.325 ± 1.193<sup>d</sup> | 0.080 ± 0.005<sup>c</sup> | 2.047 ± 0.602<sup>d</sup> |

### Gene expression and SA metabolite accumulation analyses

**i) First scenario ‘One-time inoculation’**

### Transcriptional regulation of genes involved in SA signaling, and SA accumulation in mature leaves of *C. sinensis*

Following the 7 days IAP treatment, the transcriptional regulation of *NPR1*, *NPR3*, *NPR4*, and *PR-1* and its association with endogenous SA levels were analyzed in leaves of *C*Las-infected and sham control plants (Fig 1). Quantitative RT-PCR (qRT-PCR) analysis indicated that the relative expression of *NPR1* was only significantly upregulated (*P* = 0.035) in *C*Las-infected mature leaves during the first instance of *C*Las detection (1^st^ D) compared to the sham control (Fig 1A). The relative expression of *NPR3* was significantly upregulated (1^st^ D, *P* = 0.049; 2^nd^ D, *P* = 0.045; and 4^th^ D, *P* = 0.007) during each of the three instances when *C*Las was detected in infected leaves as compared to non-infected samples (Fig 1B). However, the relative expression of *NPR4* was significantly upregulated (*P* = 0.042) in *C*Las-infected leaves only during the first instance of detection (1^st^ D), and its temporal expression pattern was downregulated (*P* < 0.05) during the second instance of *C*Las detection (2^nd^ D) compared to other time-points (Fig 1C). The NPR-regulated gene, *PR-1*, was significantly upregulated during the second instance of *C*Las detection (2^nd^ D) (*P* = 0.004) in *C*Las-infected leaves compared to sham control leaves (Fig 1D), and its temporal expression was significantly upregulated (*P* < 0.05) during the second (2^nd^ D) and fourth (4^th^ D) instances of *C*Las detection compared to the initial time when *C*Las was detected in this treatment.

**Fig 1.**
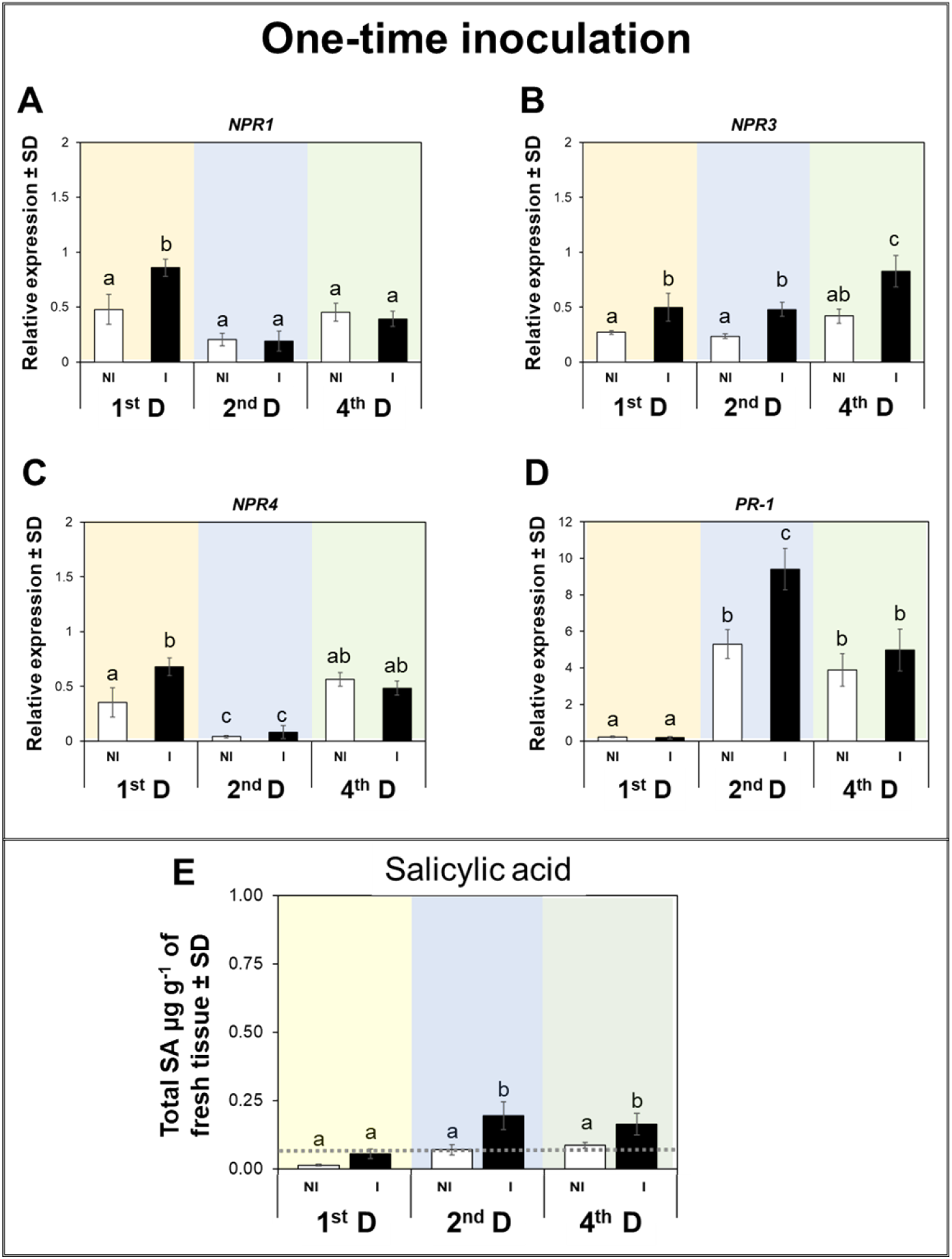
SA signaling and accumulation after ‘One-time inoculation’. CLas-infection induces changes in expression patterns of *NPR1*, *NPR3*, *NPR4*, and *PR-1*, as well as, SA accumulation in mature leaves of *C. sinensis*. Expression patterns of genes involved in SA signaling (A-D), SA accumulation in mature leaves (E). Expression levels were normalized using two references genes, *β-Actin* and *Elongation factor 1α*. Data represent mean ± standard deviation (SD) of six biological replicates per treatment. The gray dotted line denotes the level of SA accumulation at time zero in uninfected citrus plants. White bars represent plants exposed to Non-infected (NI) *D. citri*, while black bars denote plants exposed to *C*Las-infected (I) *D. citri*.

The phytohormone SA binds to NPR proteins, modulating the transcription of the *PR-1* defense gene; significantly higher levels of SA were accumulated in *C*Las-infected leaves than in control leaves during the second (2^nd^ D, *P* = 0.006) and fourth (4^th^ D, *P* = 0.049) instances of detection compared to the initial time when *C*Las was detected in this treatment (Fig 1E).

### Transcriptional regulation of genes involved in SA modifications and SA metabolite accumulation in mature leaves of *C. sinensis*

qRT-PCR analysis indicated that *BSMT,* required for the methylation of SA into MeSA, was significantly upregulated in *C*Las-infected mature leaves during the second (2^nd^ D, *P* = 0.039) and fourth (4^th^ D, *P* = 0.049) instances of *C*Las detection compared to the sham control (Fig 2A). However, *MES1*, requiring esterification of MeSA into SA (active defense hormone), did not exhibit changes in gene expression between *C*Las-infected and *C*Las-free leaves (Fig 2B). *DMR6,* involved in hydroxylation of SA into 2,3- and 2,5-DHBA, was significantly upregulated in *C*Las-infected leaves during the second (2^nd^ D, *P* = 0.006) and fourth (4^th^ D, *P* = 0.008) instances of *C*Las detection compared to *C*Las-free leaves (Fig 2C).

**Fig 2.**
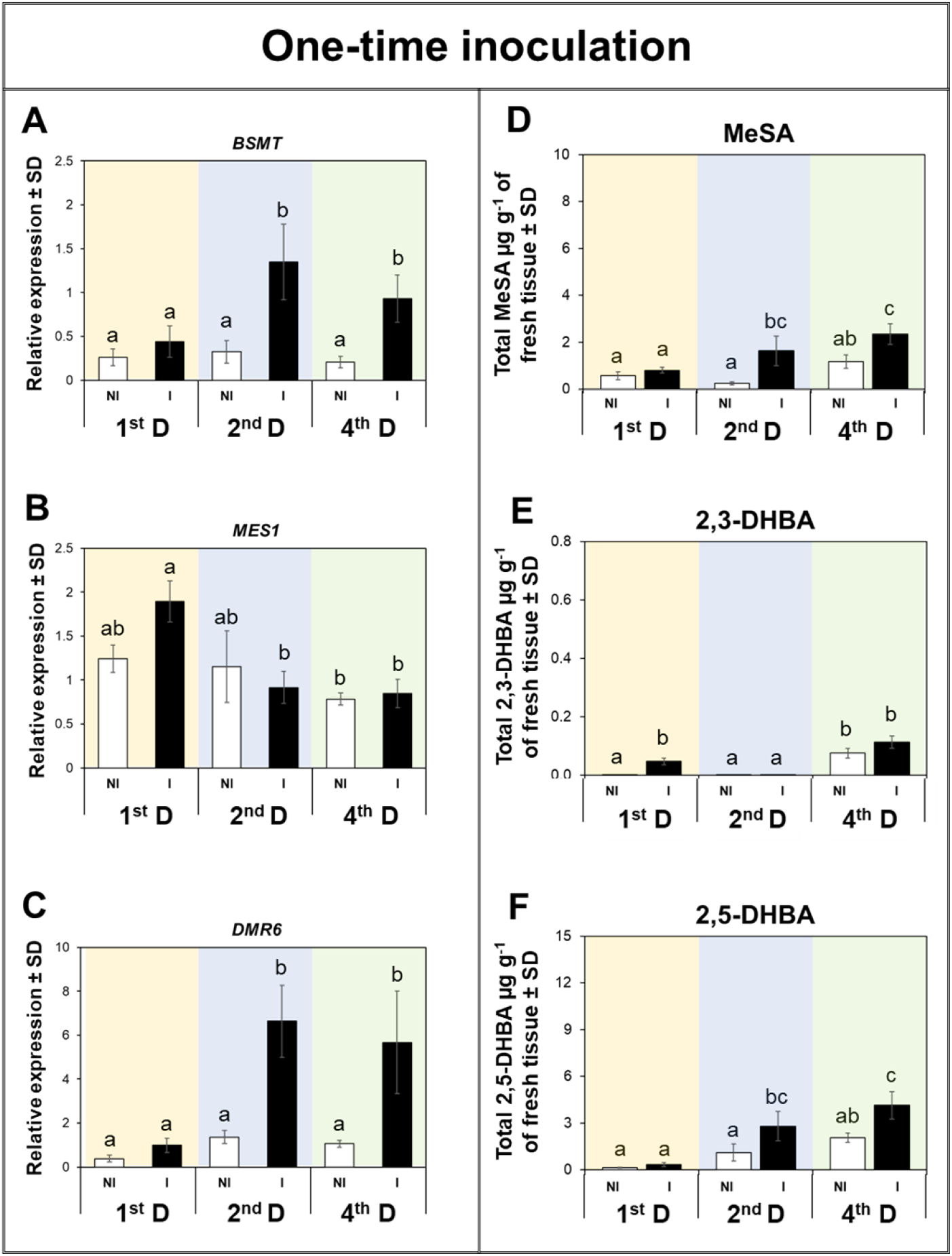
SA modifications and metabolite accumulation after ‘One-time inoculation’. CLas-infection induces changes in expression pattern of *BSMT*, *MES1*, and *DMR6*, as well as, SA metabolite accumulation in mature leaves of *C. sinensis*. Expression patterns of genes involved in SA signaling (A-C), SA metabolite accumulation in mature leaves (D-F). Expression levels were normalized using two references genes, *β-Actin* and *Elongation factor 1α*. Data represent mean ± standard deviation (SD) of six biological replicates per treatment. White bars represent plants exposed to Non-infected (NI) *D. citri*, while black bars denote plants exposed to *C*Las-infected (I) *D. citri*.

MeSA was highly accumulated in *C*Las-infected leaves at the second (2^nd^ D, *P* = 0.015) and fourth (4^th^ D, *P* = 0.032) instances of *C*Las detection. The hydroxylated molecules of SA, 2,3- and 2,5-DHBA, exhibited a different temporal accumulation; the level of 2,3-DHBA was only significantly (*P* = 0.02) higher in *C*Las-infected than control leaves during the first (1^st^ D) instance of *C*Las detection (Fig 2E). However, 2,5-DHBA was significantly accumulated in *C*Las-infected mature leaves during the second (2^nd^ D, *P* = 0.049) and fourth (4^th^ D, *P* = 0.047) instances of *C*Las detection compared to uninfected samples (Fig 2F).

### Second scenario: ‘pulsed inoculations’

qRT-PCR analysis of leaves from *C. sinensis* trees exposed to repeated (monthly) pulses of 7 days IAP revealed that relative expression of *NPR1* was significantly downregulated in *C*Las-infected leaves during the second (2^nd^ D, *P* = 0.043) and fourth (4^th^ D, *P* = 0.05) instances of *C*Las detection compared with their respective controls. Also, there was a significant reduction in the expression of *NPR1* over time in both uninfected and infected mature leaves (*P* < 0.05, Fig 3A). Expression of *NPR3* was significantly downregulated in *C*Las-infected leaves at the fourth (4^th^ D, *P* = 0.045) instance of *C*Las detection compared to the respective control (Fig 3B). However, expression of *NPR4* was upregulated in uninfected leaves during the second (2^nd^ D, *P* = 0.032) instance of *C*Las detection relative to *C*Las-infected leaves (Fig 3C). Overall, expression of all *NPR* genes tended to decrease over time in mature leaves as a response to pulses of 7-day IAPs by *C*Las-infected psyllids. Expression of *PR-1* was only upregulated in *C*Las-infected mature leaves during the second (2^nd^ D, *P* = 0.003) instance of *C*Las detection compared to its control.

**Fig 3.**
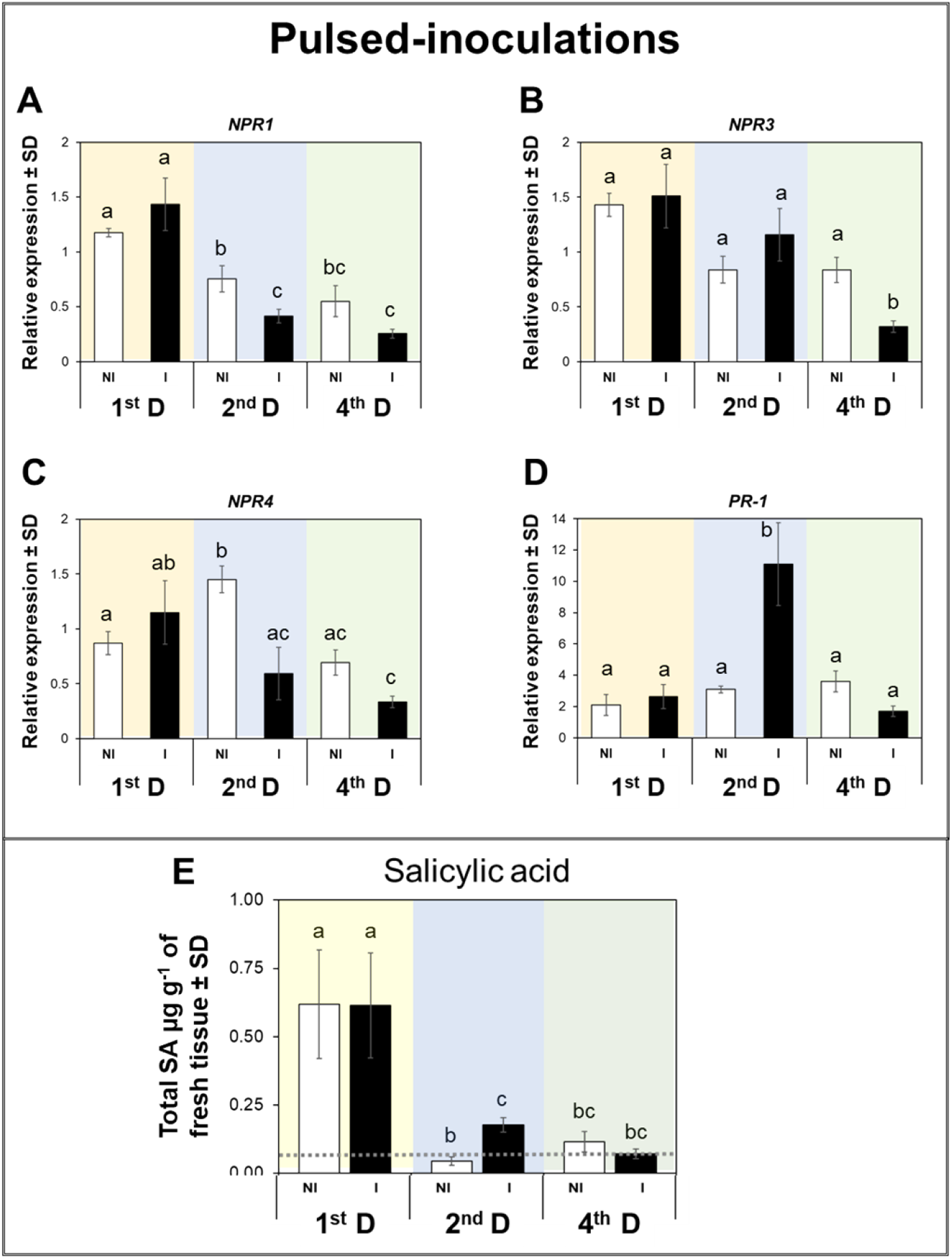
SA signaling and accumulation after ‘Pulsed-inoculations’. CLas-infection induces changes in expression pattern of *NPR1*, *NPR3*, *NPR4*, and *PR-1*, as well as, SA accumulation in mature leaves of *C. sinensis*. Expression patterns of genes involved in SA signaling (A-D), SA accumulation in mature leaves (E). Expression levels were normalized using two references genes, *β-Actin* and *Elongation factor 1α*. Data represent mean ± standard deviation (SD) of six biological replicates per treatment. The gray dotted line denotes the level of SA accumulation at time zero in uninfected citrus plants. White bars represent plants exposed to Non-infected (NI) *D. citri*, while black bars denote plants exposed to *C*Las-infected (I) *D. citri*.

Measurements of SA dynamics over time in trees that received pulses of 7-day IAPs revealed that [SA] was only significantly accumulated in *C*Las-infected mature leaves during the second (2^nd^ D, *P* = 0.049) instance of *C*Las detection compared to uninfected leaves. When temporal accumulation of SA was analyzed, we found that [SA] increased during the first (1^st^ D) instance of *C*Las detection in both uninfected and infected leaves. Thereafter, there was a significant decline of [SA] in leaves observed during the second (2^nd^ D, *P* < 0.05) and fourth (4^th^ D, *P* < 0.05) instances of *C*Las detection (Fig 3E).

### Transcriptional regulation of genes involved in SA modifications and SA metabolite accumulation in mature leaves of *C. sinensis*

The expression of *BSMT* was downregulated (*P* = 0.049) during the first (1^st^ D time) instance of *C*Las detection in *C*Las-infected leaves as compared to that observed in its control (Fig 4A). *BSMT* did exhibit significant changes in expression among sampling points over time (Fig 4A). Expression of *MES1* did not differ between *C*Las-infected and uninfected leaves (Fig 4B) at any of the sampling points; however, its expression decreased over time and was significantly (*P* < 0.05) lower during the fourth (4^th^ D) instance of *C*Las detection in *C*Las-infected than uninfected leaves. Conversely, *DMR6* was significantly downregulated (*P* = 0.045) in *C*Las-infected leaves during fourth (4^th^ D) instance of *C*Las detection compared to that observed in uninfected leaves (Fig 4C).

**Fig 4.**
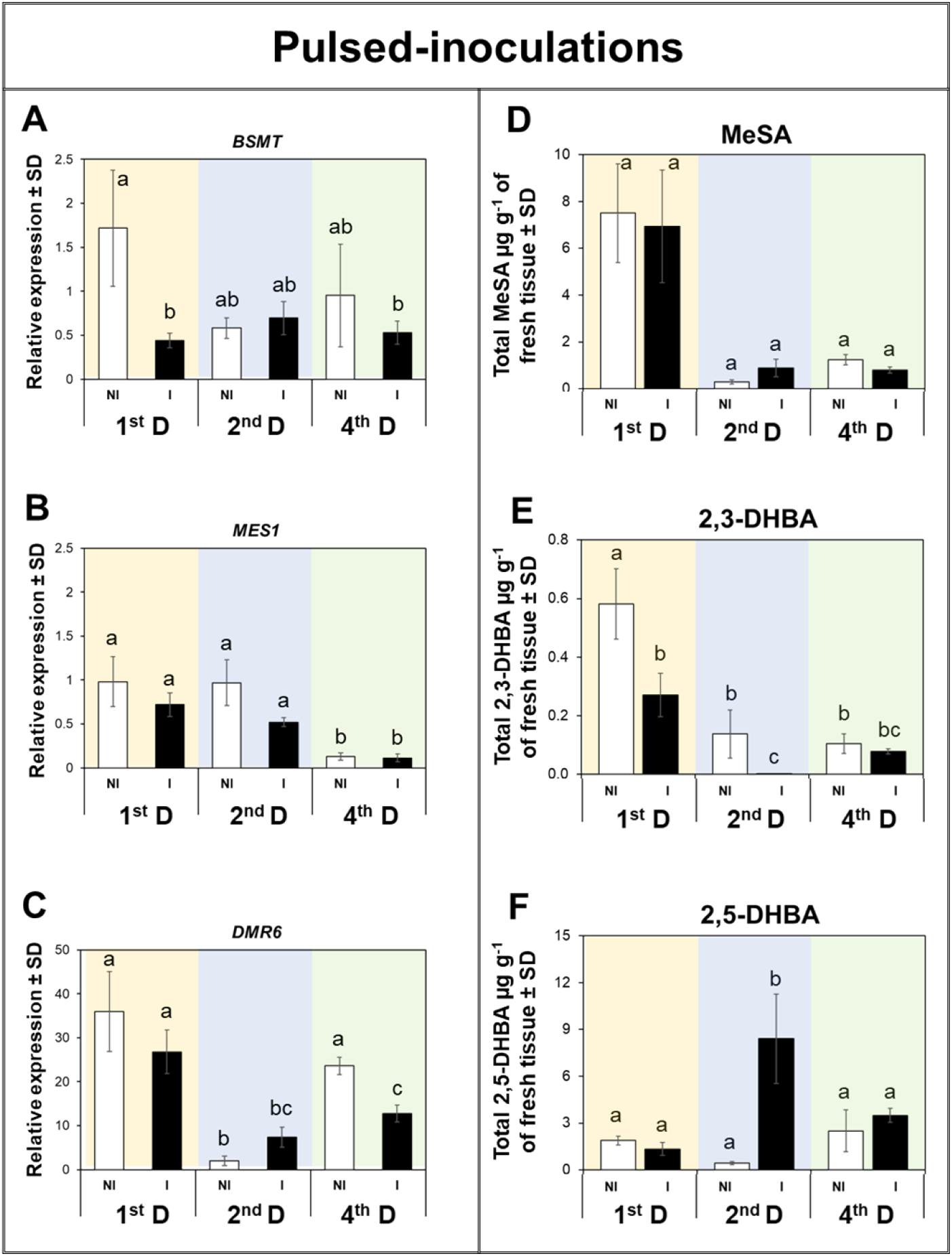
SA modifications and metabolite accumulation after ‘Pulsed-inoculations’. *C*Las-infection induces changes in expression pattern of *BSMT*, *MES1*, and *DMR6*, as well as, SA metabolite accumulation in mature leaves of *C. sinensis*. Expression patterns of genes involved in SA signaling (A-C), SA metabolite accumulation in mature leaves (D-F). Expression levels were normalized using two references genes, *β-Actin* and *Elongation factor 1α*. Data represent mean ± standard deviation (SD) of six biological replicates per treatment. White bars represent plants exposed to Non-infected (NI) *D. citri*, while black bars denote plants exposed to *C*Las-infected (I) *D. citri*.

In addition to transcriptional analysis, we examined the temporal accumulation of SA metabolites. [MeSA] did not differ significantly (*P* > 0.05) between *C*Las-infected and uninfected leaves during the specific time points when *C*Las was detected. However, there were significant (*P* < 0.05) changes in [MeSA] over time with a greater accumulation during the first (1^st^ D) instance *of* CLas detection compared to the other two time points when the pathogen was detected (Fig 4D). The [2,3-DHBA] was significantly lower in *C*Las-infected than control leaves during the first (1^st^ D, *P* = 0.045) and second (2^nd^ D, *P* = 0.041) instances of *C*Las detection. The [2,5-DHBA] was significantly higher in *C*Las-infected than uninfected leaves during the second (2^nd^ D, *P* = 0.0001) instance of *C*Las detection (Fig 4F).

### Third scenario: ‘continuous inoculations’

The transcriptional regulation of *C. sinensis* exposed to a continuously reproducing population of *D. citri* revealed that *NPR1* expression did not differ between *C*Las-infected and uninfected leaves at any of the time points when *C*Las was detected (Fig 5A). However, expression of *NPR1* was significantly (*P* < 0.05) lower during the second (2^nd^ D) and fourth (4^th^ D) instance of *C*Las infection as compared its expression when *C*Las was detected for the first time (Fig 5A). The expression of *NPR3* was upregulated in *C*Las-infected leaves during the first (1^st^ D, *P* = 0.046) instance of *C*Las detection compared with that in its respective uninfected control (Fig 5B), and similarly to *NPR1*, its expression was downregulated during the second (2^nd^ D) and fourth (4^th^ D) instance of *C*Las detection in *C*Las-infected as compared with uninfected leaves (Fig 5B). In the case *NPR4*, there were no statistical differences between infected and uninfected plants exposed to reproducing populations of *D. citri* (Fig 5C). Expression of *NPR4* was significantly (*P* < 0.05) lower during the fourth (4^th^ D) instance of *C*Las detection as compared its expression when *C*Las was detected for the first time (Fig 5C). Expression of *PR-1* was significantly upregulated (*P* < 0.001) in uninfected than *C*Las-infected leaves during the second (2^nd^ D) instance of *C*Las detection (Fig 5D).

**Fig 5.**
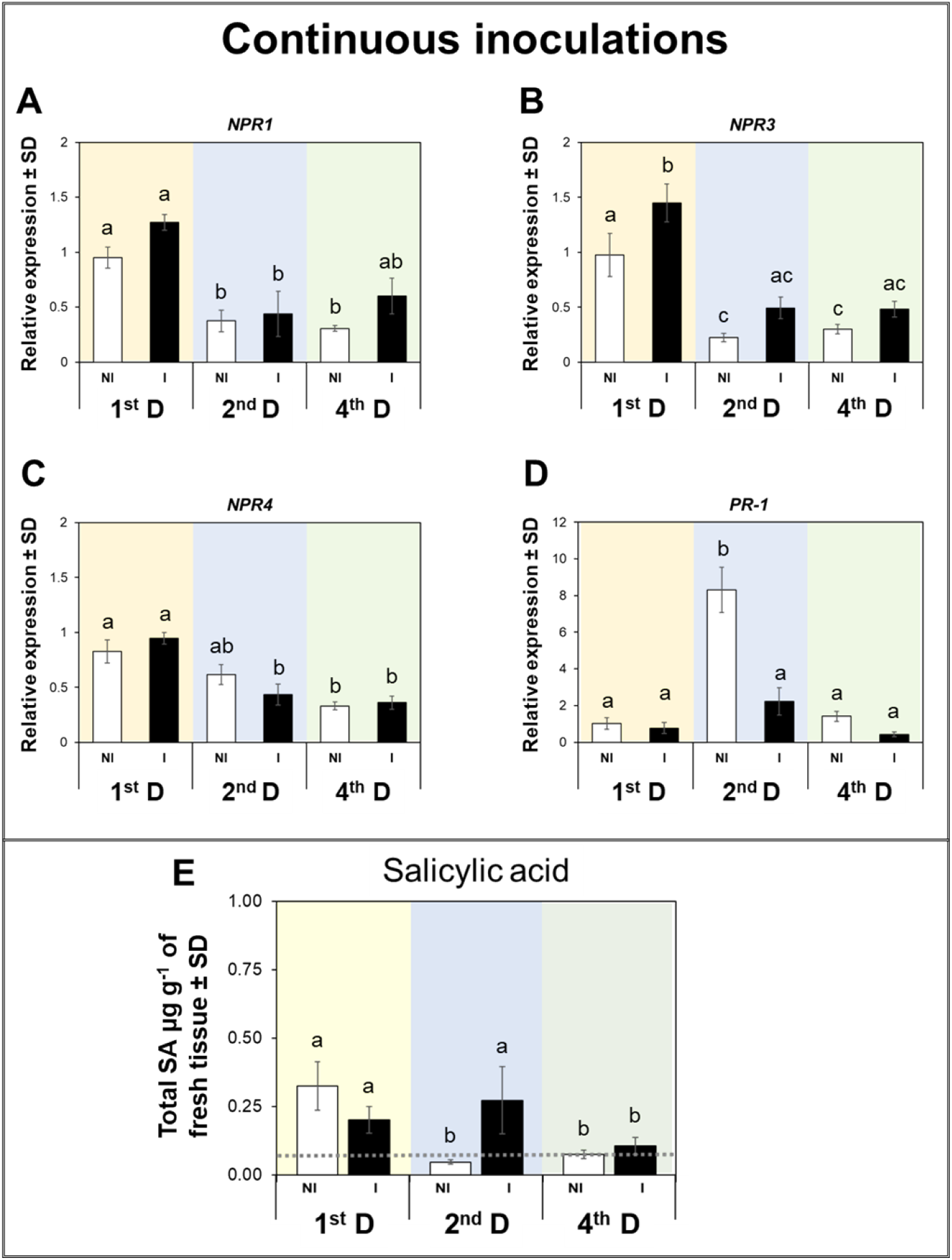
Signaling of SA and accumulation after ‘Continuous-inoculations’. CLas-infection induces changes in expression pattern of *NPR1*, *NPR3*, *NPR4*, and *PR-1*, as well as, SA accumulation in mature leaves of *C. sinensis*. Expression patterns of genes involved in SA signaling (A-D), SA accumulation in mature leaves (E). Expression levels were normalized using two references genes, *β-Actin* and *Elongation factor 1α*. Data represent mean ± standard deviation (SD) of six biological replicates per treatment. White bars represent plants exposed to Non-infected (NI) *D. citri*, while black bars denote plants exposed to *C*Las-infected (I) *D. citri*.

The [SA] was higher in *C*Las-infected plants exposed to continuous re-inoculation and herbivory during the second (2^nd^ D time) instance of *C*Las infection compared to the complementary control plants that were exposed to uninfected insects. However, the [SA] was significantly (*P* > 0.05) lower during the fourth (4^th^ D) instance of *C*Las detection compared with [SA] recorded in both infected and uninfected leaves when *C*Las was detected for the first time (Fig 5E).

Expression of *BMST* was significantly higher in *C*Las-infected than uninfected leaves during the second (2^nd^ D) instance of *C*Las infection among plants that were continuously exposed to reproducing *D. citri* (*P* = 0.002, Fig 6A). The expression pattern of *MES1* did not differ between *C*Las-infected and uninfected samples; however, its expression was significantly upregulated (*P* < 0.05) during the second (2^nd^ D) instance of *C*Las detection in both treatments as compared to that observed during the other sampling points (Fig 6B). The expression of the transcript involved in SA hydroxylation, *DMR6*, fluctuated over time with significantly (*P* = 0.007) higher expression during the second (2^nd^ D) instance of *C*Las detection in *C*Las-infected than uninfected leaves and lower overall expression during the fourth (4^th^ D) instance of *C*Las detection in both treatments (*P* < 0.05, Fig 6C).

**Fig 6.**
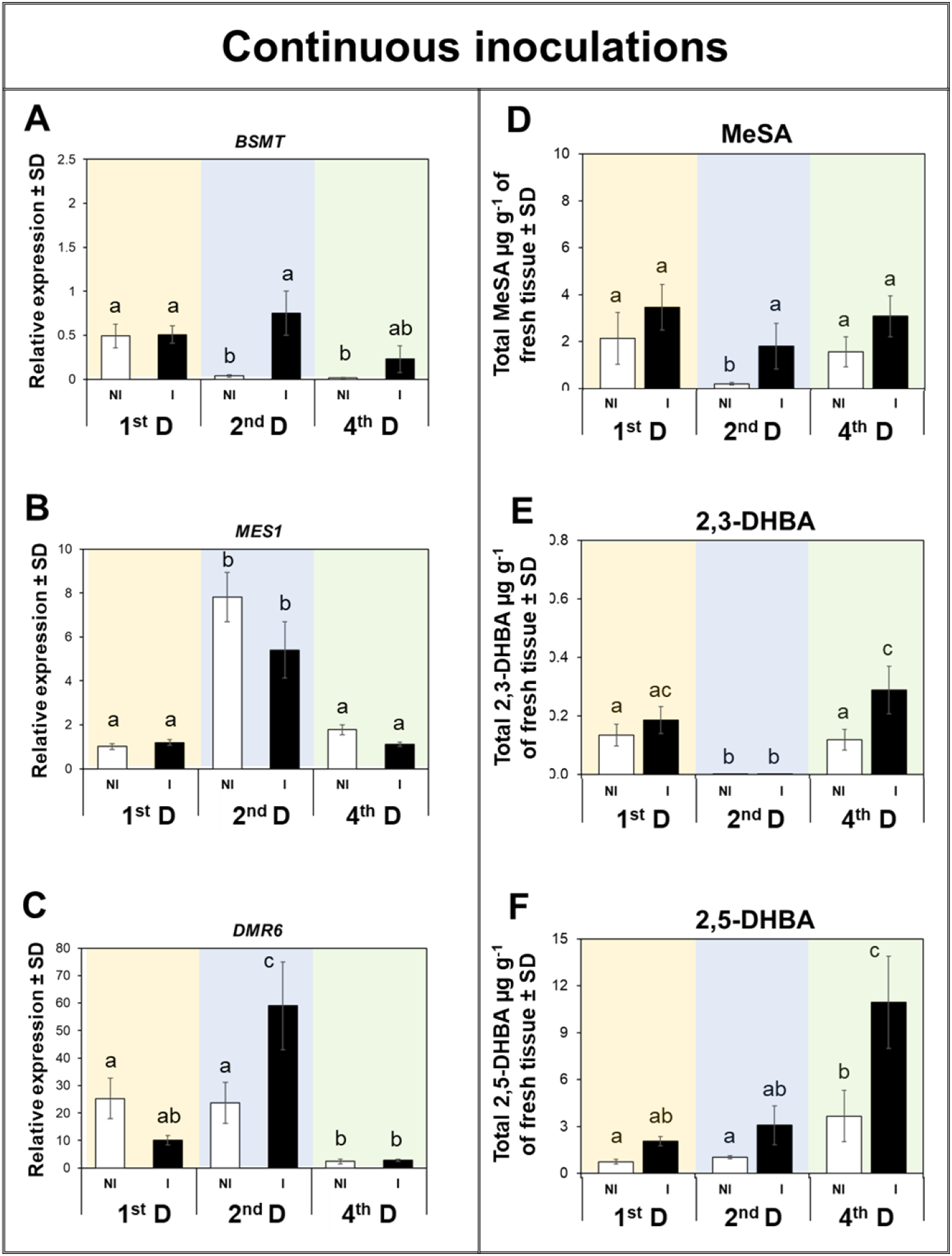
SA modifications and metabolite accumulation after ‘Continuous-inoculations’. CLas-infection induces changes in expression pattern of *BSMT*, *MES1*, and *DMR6*, as well as, SA metabolite accumulation in mature leaves of *C. sinensis*. Expression patterns of genes involved in SA signaling (A-C), SA metabolite accumulation in mature leaves (D-F). Expression levels were normalized using two references genes, *β-Actin* and *Elongation factor 1α*. Data represent mean ± standard deviation (SD) of six biological replicates per treatment. Statistical analysis was performed with repeated measures ANOVA and Tukey post-hoc tests. Different letters indicate significant differences between the samples (*P* < 0.05). White bars represent plants exposed to Non-infected (NI) *D. citri*, while black bars denote plants exposed to *C*Las-infected (I) *D. citri*.

The [MeSA] was significantly higher in *C*Las infected than uninfected leaves when both treatments were exposed to continuous herbivory only during the second (2^nd^ D, *P* < 0.05) instance of *C*Las detection (Fig 6D). Of the hydroxylated [SA] metabolites, the [2,3-DHBA] was significantly higher (*P* = 0.024) in leaves of plants exposed to *C*Las-infected insects than in plants exposed to uninfected insects during the fourth (4^th^ D) instance of *C*Las detection (Fig 6E). However, [2,5-DHBA] was only significantly higher in *C*Las-infected than uninfected leaves during the fourth (4^th^ D) instance of *C*Las detection (*P* = 0.001), and there was a trend of progressively increasing 2,5-DHBA in leaves exposed to both infected and uninfected insects over time (Fig 6F).

### Comparative analyses of SA and its analytes in response to the interaction of insect herbivory and pathogen infection

When the accumulated amount of SA and its metabolites were compared between all three scenarios investigated, there was a statistically significant increase of [SA] in *C*Las-infected leaves of plants exposed to infected *D. citri* as compared with the uninfected sham treatment in the OI scenario (Fig 7), suggesting a *C*Las-induced SA response. However, there was greater overall accumulation of SA in mature leaves from trees under the PI and CI scenarios than that accumulating in the OI scenario. Furthermore, when comparing between PI and CI scenarios, a difference in accumulation of SA between the *C*Las infection treatment and sham control was not apparent (Fig 7A).

**Fig 7.**
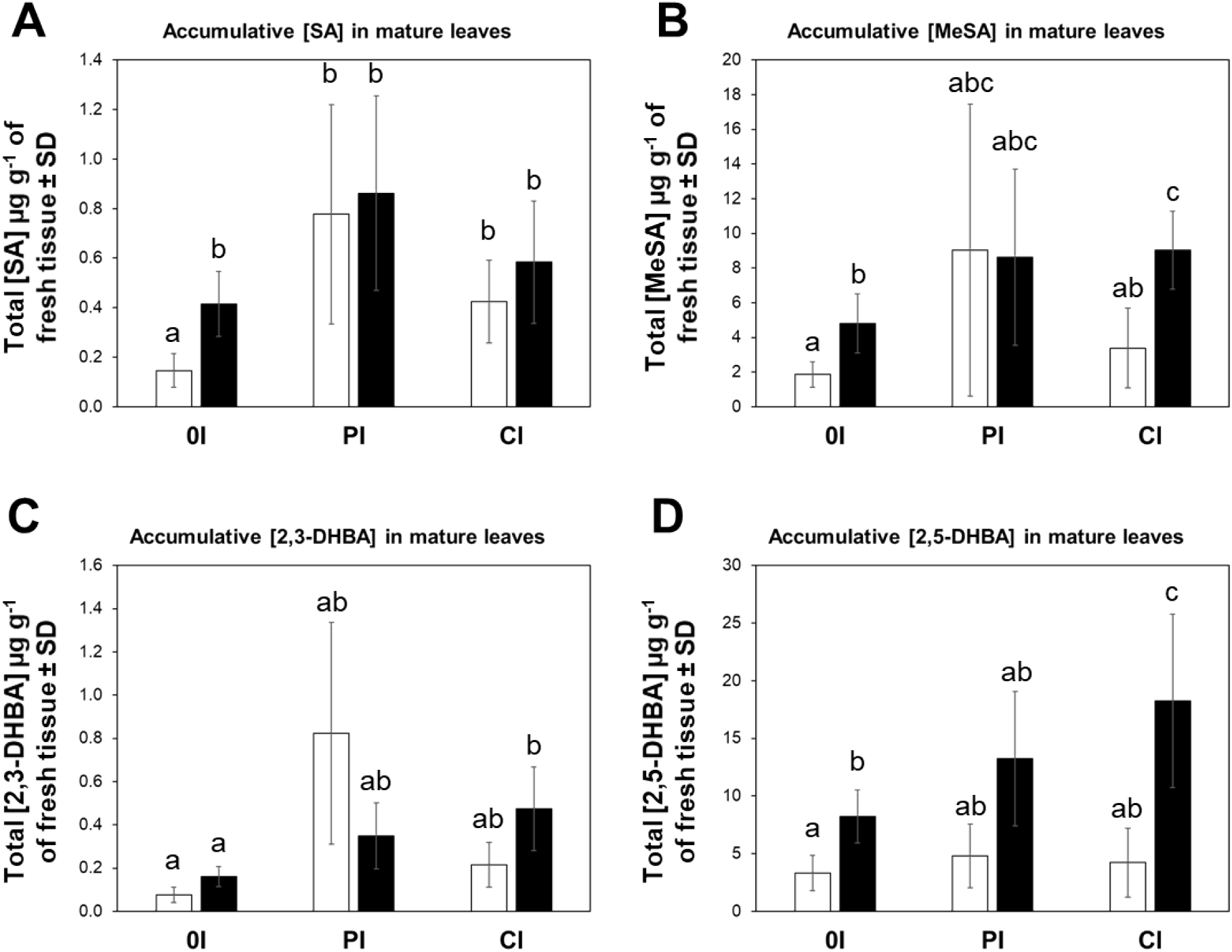
Cumulative SA and metabolite concentrations among different scenarios of a vector-host-pathogen interaction in citrus. Data represent mean ± standard deviation (SD) of six biological replicates per treatment. Statistical analysis was performed with ANOVA with Tukey post-hoc tests. Different letters indicate significant differences between the samples (*P* < 0.05). White bars represent citrus plants exposed to uninfected psyllids, and black bars denote plants challenged with *C*Las-infected psyllids, (OI = One-time inoculation, PI = Pulsed inoculations and CI = continuous inoculations).

## Discussion

Plants perceive microbial and insect molecules as danger signals and mount defense against invasions. Studies investigating plant interactions with phytopathogens have established that plants have a layered innate immune system which responds to different microbial elicitors [5], and early signaling events are similar to those induced by insects [34–37]. The current investigation attempted to both disentangle the contributing effects of insect herbivory and pathogen infection on host plant response, as well as, gain insights into their interactive effects on disease progression. We inflicted insect and pathogen activity on citrus hosts with manipulative treatments that were designed to mimic the types of infestations/infections that citrus growers experience when cultivating citrus in the face of Huanglongbing (HLB). *C. sinensis* plants were challenged with three putative scenarios of vector-host-pathogen interactions [one time (OI), pulsed inoculation PI, and constant inoculation (CI] to explore transcriptional and metabolic modulation of SA-dependent defense responses in the host after varying frequencies of herbivore infestation and/or *C*Las inoculation. Moreover, the investigation was longitudinal, which allowed sampling of gene expression and metabolite accumulation over the course of disease progression.

Progression of pathogen in the host after trees were exposed to various frequencies of vector IAP (with or without pathogen present) was measured by monitoring *C*Las DNA. Simultaneously, we measured defense responses in *C. sinensis*, which may combat pathogenesis induced by *C*Las. Surprisingly, our results indicate that *C. sinensis* trees challenged by *C*Las-infected insects for only a 7d IAP exhibited similar temporal dynamics of *C*Las titer to trees that were continuously challenged with infected *D. citri* for over 1 yr. In contrast, trees exposed to many (12-14), repeated 7d IAP challenges (pulsed-inoculations) by infected *D. citri* exhibited the lowest *C*Las titer, suggesting putative short-term plant defense responses to *C*Las (HLB progression, infection). We hypothesized that SA-dependent defense responses were active in these trees to reduce or ‘control’ *C*Las colonization.

Our results suggest that herbivory played an important role in inducing synthesis and accumulation of SA in mature leaves of *C. sinensis*, whether or not those trees are coincidentally infected with *C*Las. Accumulation of SA and the SA-signaling pathway play a critical role in plant defenses against pathogens [20] and in response to phloem-feeding herbivores [21–23]. SA regulates PTI and ETI defense mechanisms which leads to activation of systemic acquired resistance (SAR). We postulate that it might be eventually possible to manipulate natural defense and immune responses of *Citrus* to *C*Las and/or *D. citri* with sufficient understanding of the genes involved in SA-dependent immune response (*NPR1*, *NPR3*, *NPR4*, and *PR-1*) and genes that modulate downstream metabolic modifications (*BMST*, *MES1*, and *DMR6*).

Previously, a gene involved in the hydroxylation of SA,*SA hydroxylase*, was identified and characterized in *C*Las; infected plants exhibited significantly lower SA accumulation in leaves than uninfected plants 8 months after graft inoculation [38]. However, a similar reduction in SA was not observed in other investigations of the *C*Las-*C. sinensis* interaction. For example, Lu, Zhang (39) showed that symptomatic and PCR-positive leaves collected 3-4 months after graft-inoculation exhibited higher SA accumulation than uninfected trees. Similar results were detected after 6.4 months in young and mature leaves from trees previously graft-inoculated with *C*Las [40]. In the first scenario we investigated, (OI) - trees without insects after initial IAP, significantly more SA accumulated in leaves from infected than uninfected trees at 9 and 12 months (Fig 1E). This outcome suggests that *C*Las did not evade SA-mediated immune responses via activity of its encoded SA hydroxylase within the context of this scenario where pathogen occurred *in planta* in the absence of concurrent insect herbivory/re-inoculation.

In addition to upstream regulation of SA, this hormone is also modulated by downstream metabolic modifications, including glycosylation, methylation, amino acid conjugation, and hydroxylation [25, 27]. In this investigation, production of the methylated (MeSA) and hydroxylated SA-metabolites (2,3- and 2,5-DHBAs) was measured longitudinally (Figs 2, 4 and 6) and their cumulative amounts were determined for the duration of the experiment (Fig 7 B-D). MeSA serves multiple functions, including triggering of SAR in response to microbial pathogens as shown with Arabidopsis, tobacco, and potato [41–44]. MeSA also functions as an indirect defense cue by attracting natural enemies in response to insect feeding [45–47]; e.g. MeSA attracts *D. citri* to citrus trees [48]. We measured substantial accumulation of MeSA in *C*Las-infected leaves (OI and PI). When *C*Las-infected vector IAP occurred in repeated pulses (the PI scenario), the titer of *C*Las in infected plants was the lowest of all three infection scenarios investigated (Table 1). In trees subjected to this IAP scenario, the concentration of MeSA varied over time and no difference in concentration was observed between trees injured by *C*Las-infected and uninfected insects (Fig 7B). In contrast, when plants were subjected to a one time IAP (OI) or a continuously breeding population of vectors (CI), the [MeSA] was substantially higher in leaves from trees that were also challenged with the pathogen than in the sham controls that received only insect injury in the absence of *C*Las. These results are congruent with previous observations showing that *C*Las-infected field trees are attractive to *D. citri*, which is, in part, explained by MeSA serving an attractant cue or the vector (Mann et al. 2012).

Although the specific roles played by the hydroxylated SA molecules, 2,3- and 2,5-DHBA, in plant immune defense have not yet been clearly elucidated, studies suggest that exogenous applications of 2,3-DHBA in Arabidopsis induced weak *PR-1* expression [49]. Furthermore, 2,5-DHBA induced the synthesis of a different set of PR proteins in tomato [50]. We found no difference in 2,3-DHBA accumulation between trees that were subjected to sham (uninfected) IAPs versus *C*Las-infected IAPs, which resulted in eventual tree infection. However, more 2,3-DHBA accumulated in *C*Las-infected leaves that were subjected to a continuously breeding population of vectors (CI scenario) than in *C*Las-infected leaves from trees that were only injured by vectors during an initial one-time 7-d IAP (OI scenario, Fig 7C). In the case of 2,5-DHBA, it was highly accumulated in *C*Las-infected leaves (both OI and CI scenarios) compared to their respective sham controls, while the pulsed inoculation (PI scenario) caused intermediate accumulation with no difference observed between *C*Las-infected and uninfected (sham IAP) trees (Fig 7D). Based on our results, it appears that *C*Las infection induces the accumulation of 2,5-DHBA in *C. sinensis*; however, the effect of this analyte within the context of plant immune response is still unclear. The elucidation of this function was beyond the scope of our current study, but the role(s) of these hydroxylated SA molecules in *Citrus* should be investigated further.

### Gene expression and its association with SA and related metabolites

We propose a working model of SA-modulated defense mechanisms in *C. sinensis* (S1 Fig) that examines the expression patterns of *NPR1*, *NPR3*, *NPR4*, *PR-1*, *BMST*, *MES1*, and *DMR6*, as well as, the accumulation of SA and its hydroxylated and methylated analytes based on the three scenarios of vector interactions with the host and pathogen (OI, PI, and CI).

#### One-time inoculation scenario

Using Arabidopsis and other plant species as models, it has been proposed that overexpression of *NPR1* induces expression of disease response genes, such as *PR-1* [51–53]. In our first scenario, *C*Las-infected samples exhibited a transient upregulation of *NPR1* in mature leaves only at 120 days after initial IAP and this coincided with the initial detection of *C*Las in the leaf phloem. However, upregulation of *NPR1* was not associated with a similar *PR-1* expression pattern as expected; *PR-1* was upregulated in *C*Las-infected leaves much later after initial IAP at 270- and 330- days, which was also much later than the initial detection of *C*Las titer in phloem of infected trees (day 120). Furthermore, this *PR-1* expression pattern was associated with high accumulation of SA in *C*Las-infected samples and reduced expression of *NPR4*. Recently, it was shown in Arabidopsis that the transcription factors NPR3/NPR4 interact with TGA2/TGA5/TGA6 to downregulate the expression of genes involved in plant defenses when plants are not infected and exhibit low SA concentration. Also, a gain-of-function mutation, *npr4-4D*, leads to a SA-insensitive mutant protein that represses SA-inducible defense genes [54]. Consequently, the expression pattern of *PR-1* is congruent with low expression of *NPR4* in association with SA accumulation in *C*Las-infected leaves. Therefore, we suggest that infection of *C. sinensis* with *C*Las activates SA-dependent defense responses even in the absence of phloem feeding injury caused by the vector. However, characterizing how the pathogen induces this SA response in *C. sinensis* would likely require investigations employing gain/loss of function mutations, which will only be possible once this fastidious bacterium is reliably cultured [55–59].

#### Pulsed-inoculations scenario

For this inoculation scenario, trees were challenged by monthly ‘pulses’ of 7 d IAP inflicted by *C*Las-infected adult psyllids. To our surprise, these trees appeared to exhibit a much higher degree of ‘tolerance’ against *C*Las colonization, given that the lowest pathogen titers were measured in these trees when comparing across the three infection scenarios that we examined. In the pulsed inoculation scenario, SA-dependent immune responses and associated accumulation of SA and its metabolites were both substantial at 120 days after initial IAP, but then progressively decreased over time (Figs 4 and 5). We postulate that repeated exposure to insect injury caused sufficient accumulation of SA in *C. sinensis* to inhibit normal *C*Las colonization. Furthermore, we observed high expression of *PR-1* coincident with downregulation of *NPR1* and *NPR4* in trees exposed to *C*Las-infected insects by 270 days after initial IAP (Fig 3) even though no bacterial DNA from the pathogen was detected in leaf phloem at this point in the experiment (Table 1).

#### Continuous inoculations scenario

In this scenario, trees were challenged by the possibility of continuous *C*Las re-infection and constant injury inflicted by a reproducing population of *D. citri*. In general, the pattern of *NPR* transcripts differed from the OI scenario in that it decreased over time in both uninfected (sham inoculation receiving insect injury only) and *C*Las-infected trees; this downregulation of *NPR* response was also associated with lower SA accumulation than in the OI scenario. Specifically, trees that were exposed to continuous feeding by *D. citri* initially accumulated SA (120 days after initial IAP), but SA production decreased significantly at later time-points (270- and 330- days after initial IAP). Injury to *C. sinensis* by *D. citri* feeding for brief intervals (≤14 days) and in the absence of *C*Las infection induces expression of *NPR1* and *PR-1*, but is not associated with changes in SA accumulation compared to control plants (without *D. citri*) [23]. In contrast, prolonged (≥ 150) feeding injury by uninfected *D. citri* stimulates accumulation of SA, yet no changes in expression of *NPR1* and *PR-1* were detected in these trees [23]. Collectively, these results indicate that psyllid feeding injury stimulates immune defenses in *C. sinensis* soon after feeding injury is initiated which can result in accumulation of SA, but prolonged feeding turns off SA synthesis and/or increases expression of genes involved in SA metabolism (*BMST* and *DMR6*), resulting in accumulation of SA metabolites (MeSA, 2,3-DHBA, and 2,5-DHBA) (Fig 7).

Our investigation describes gene expression associated with SA-dependent immune responses, as well as, resultant accumulation of SA and its metabolites in *C. sinensis* after various scenarios simulating infection of *Citrus* with the pathogen causing HLB. The investigation considered the interaction between vector and pathogen on host responses and varied duration of exposure to these two stressors. Based on our findings, we postulate that SA-dependent defense responses were activated in *Citrus* trees exposed to pulses of *C*Las inoculation at least until the initial instance of *C*Las detection in phloem. SA responses are of particular relevance to understanding HLB disease progression in sweet oranges, given that increased callose and phloem protein 2 deposition occurs in sieve elements of citrus trees expressing HLB symptoms [60, 61]. A recent investigation by Deng, Achor (4), showed that ‘Bearss’ lemon and ‘LB8-9’ Sugar Belle^®^ mandarin were more tolerant to *C*Las infection than most commercial citrus varieties, which was associated with lower levels of phloem disruption and greater phloem regeneration than in HLB-susceptible sweet orange. It is unclear whether or not SA was associated with this phenomenon; thus, it is imperative to understand the temporal accumulation of SA in citrus exposed to *D. citri* (vector) herbivory, as well as, *C*Las (pathogen) infection, and use this information to address fundamental questions related to symptom suppression and associated practical questions relating to developing tree therapy. Determining gene function involved in immune responses and SA modification in *Citrus* may yield novel genotypes that express tolerance against *C*Las and/or modify vector behavior to reduce inoculation.

The abnormal levels of gene expression and metabolite accumulation observed in trees challenged by prolonged feeding injury by *D. citri* appears to have a deleterious effect on the immune response of *C. sinensis*. From an immediately practical perspective, these findings support the practice of vector suppression as part of HLB management in citrus, even in areas where HLB is established endemically [23].

## Methods

### Controlled microcosm experiments

#### Plant husbandry

All experiments were performed using of 2 year old *Citrus sinensis* L. Osbeck cv Valencia grafted onto US-812 rootstocks [62]. Plants were obtained from Southern Citrus Nurseries, Dundee, Florida. The process of eliminating possible insecticide residues from plants prior to initiation of experiments and in depth growing conditions have been described previously [23]. In brief, plants were maintained at 23 ± 3 °C, 60RH, and a 16:8 h (Light: Dark) photoperiod. Plants were watered twice per week, and fertilized once per month with an alternating schedule of a 24-8-16 (Nitrogen–Phosphorus–Potassium) Miracle-Gro All Purpose Plant Food (Scotts Miracle-Gro Products, Marysville, OH) and a 10-10-10 (N–P–K) granular fertilizer (Growers Fertilizer Corp., Lake Alfred, FL).

#### Insect rearing conditions

Colonies of *C*Las-free and *C*Las-infected of *D. citri* were maintained on *C. sinensis* L. Osbeck cv Valencia trees in two separate greenhouses. Each greenhouse was maintained at 26 ± 2°C, 60-65% RH, and a 16:8 h (Light: Dark) photoperiod with a maximum photosynthetic radiation of 215 μmol s^−1^ m^−2^. The *C*Las-infection rate was determined in each population monthly by testing a sub-sample of 40 adult insects per colony (infected and uninfected) using a TaqMan qPCR assay following the protocol described by Li, Hartung (63).

#### Inoculation frequency and herbivory duration treatments

Treatments were imposed on sweet orange, *C. sinensis*, seedling trees to simulate possible interactive scenarios of pathogen infection and insect herbivory that occur citrus groves with HLB. For each scenario below, 6 replicate trees were treated with CLas-infected or sham uninfected (control) psyllids. All trees (treatments and sham controls) were maintained individually in insect-proof cages (58.4 × 58.4 × 88.9 cm). Trees of similar phenology, characterized by presence of bud breaks and/or feather flush-like structures, were exposed to various frequencies of *C*Las-inoculation access periods (IAP) and duration of insect herbivory, as follows:

i. **First scenario: ‘one-time inoculation’ (OI)**: Plants were exposed individually to groups of 20 *C*Las-infected (treatment) or CLas-free (control) *D. citri* adults (female: males; 1:1) for a 7d IAP. On day 7, all eggs laid on flush shoots and *D. citri* adults were gently removed and *C*Las-infection of adult psyllids was determined by TaqMan qPCR assay. This treatment simulated a scenario where a citrus grove is infested by a unique immigrant *D. citri* population that is eliminated by insecticide within one week under a zero-tolerance protocol.
ii. **Second scenario: ‘pulsed inoculations’ (PI)**: Individual plants (N = 6 plants) were exposed to 20 *C*Las-infected (treatment) or *C*Las-free (sham control) *D. citri* adults for a 7d IAP. Psyllids were placed on branches within mesh sleeve cages. On the seventh day following insect release onto plants, all psyllids within cages were gently removed. This process was repeated monthly for the duration of the experiment (12 pulses of inoculations). In order to determine the *C*Las-infection rate of the adult insects throughout the course of this experiment, a subset of 40 psyllids were collected and examined by TaqMan qPCR assay during each month of the experiment; the rate of infection in psyllids varied between 35-72%. This treatment simulated a scenario where a citrus grove not harboring an endemic population of psyllids is periodically infested by immigrating psyllids, which are killed off by monthly applications of insecticides with short residual control.
iii. **Third scenario:** ‘**continuous inoculations’ (CI)**: Individual plants were exposed to 20 *C*Las-infected (treatment) or *C*Las-free (sham control) *D. citri* by releasing psyllids into cages containing trees. In this treatment, insects were not removed and plants were constantly exposed to continuously reproducing *D. citri* populations. In order to estimate the *C*Las-infection rate of the reproducing *D. citri* populations within cages, groups of insects (40 / tree) were randomly collected at three specific time-points between August (2018) and February (2019). This treatment was designed to simulate a scenario where a grove is continuously infested by an endemic population *of D. citri* that is never eliminated by management practices.

Plant tissue was sampled monthly from the same 12 mature leaves on each tree. Samples were collected by harvesting discs from these leaves beginning at the leaf tip and progressing down the petiole on each sampled leaf as described in [64, 65]. Plant tissue samples were immediately flash frozen in liquid nitrogen, ground with a tissue lyser, and stored at −80°C for further analyses.

#### Genomic DNA extraction and CLas detection by TaqMan qPCR assays

To analyze the *C*Las infection rate in insects and plants, genomic DNA from individual *D. citri* adults and 20 mg of plant tissues was extracted using the DNeasy blood and tissue or DNeasy plant kits (Qiagen Inc, Valencia, CA), respectively. Both DNA extraction protocols were completed according to manufacturer’s instructions. Quantity of genomic DNA was determined in a Nanodrop 2000 Spectrophotometer (Thermo fisher Scientific, Waltham, MA), and samples were stored at −20°C until TaqMan qPCR analysis were performed.

TaqMan assays were conducted using specific TaqMan probes previously designed by Li, Hartung (63) and targeting the 16S rDNA for *C*Las, *Wingless* for *D. citri*, and *Cytochrome oxidase I* for *C. sinensis*. TaqMan qPCR reactions and conditions were carried out as previously described [65]. TaqMan reactions were performed in duplicates and positive reactions were considered for either target sequence if the cycle quantification (Cq) value was ≤ 36. For *C*Las detection analysis, ‘undetermined values’ were automatically assigned with a Cq value equal to 45 cycles.

#### RNA extraction and cDNA synthesis

Total RNA extractions were carried out using 30 mg of ground plant tissue and the RNeasy Plant mini kit (Qiagen) according to the manufacturer’s instructions. To eliminate traces of genomic DNA, 1 μg of total RNA per sample was treated with DNase I using the Turbo DNase kit (Ambion), following the manufacturer’s protocol. RNA quantity and purity were analyzed in a Nanodrop 2000 Spectrophotometer (Thermo fisher Scientific, Waltham, MA).

cDNA synthesis was executed using the Verso cDNA Synthesis kit (Thermo Fisher scientific, CA). Each reaction consisted of: 500 ng of total RNA, 5X cDNA synthesis buffer, anchored-Oligo (dT) primers, RT Enhancer, and Verso Enzyme Mix following the manufacturer’s instructions. After synthesis, samples were stored at −20°C until further analyses.

#### Selection of genes involved in SA signaling and metabolism

In order to examine the transcriptional regulation of seven genes involved in the SA-dependent defense responses on *C. sinensis*, we selected and evaluated four genes involved in the SA signaling; *Nonexpresser pathogenesis-related 1* (*NPR1*, accession number XM_006475416)*, Nonexpresser pathogenesis-related 3* (*NPR3*, accession number XM_006468378)*, Nonexpresser pathogenesis-related 4* (*NPR4*, accession number XM_025100491), and the *Pathogenesis-related 1* (*PR1,* accession number XM_006486757). Also, we examined three genes related to SA metabolism: *Methylesterase 1* (*MES1,* accession number NM_127926)*, 2-oxoglutarate (2OG) and Fe(II)-dependent oxygenase* (Accession number NM_117118) annotated and described by [66] as a *salicylic acid 3-hydroxylase* (*DMR6*), and *S-adenosyl-L-methionine-dependent methyltransferases* (*BSMT1*, Accession number NM_111981). The expression patterns of these genes in response to feeding by uninfected *D. citri* were previously described [23]. All oligonucleotide primers for RT-qPCR analysis were designed using Primer3 web [67].

#### RT-qPCR reactions, gene expression, and metabolite analyses

RT-qPCR reactions were performed using an Applied Biosystems 7500 Real-Time PCR System (Thermo Fisher Scientific). Each reaction consisted of: 10 ng of cDNA (template), 300 nM of each gene-specific primer (Table 2) and 1x of PowerUp™ SYBR® Green Master Mix; the final volume was adjusted with nuclease-free water to 20 μL. The real-time PCR program, conditions, normalization, and mathematical estimations of relative expression at different time-points were conducted according to Ibanez, Suh (23).

**Table 2.**
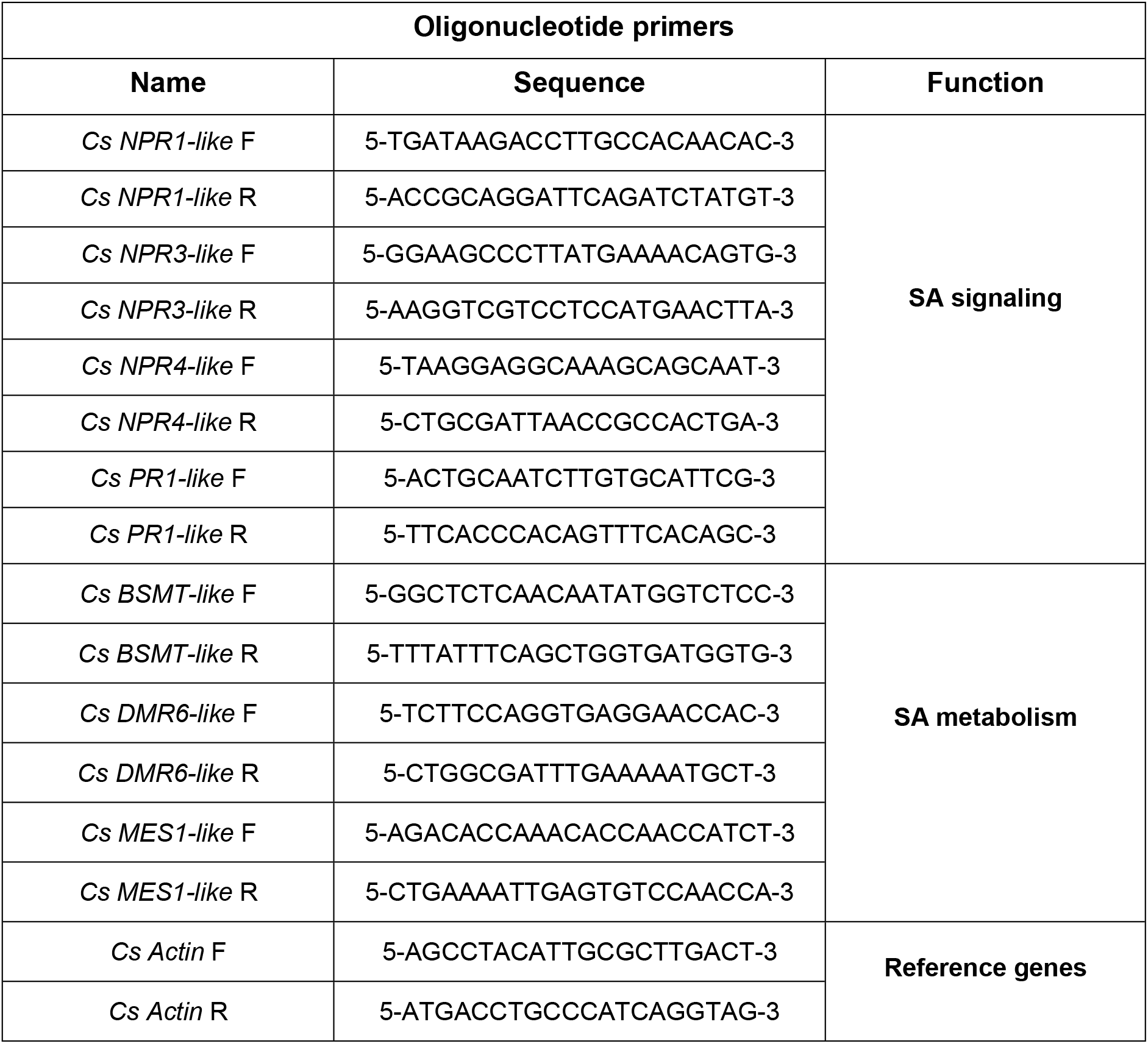
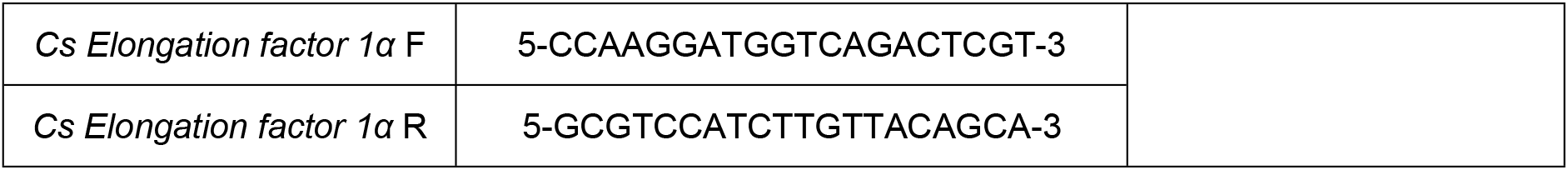
Oligonucleotide primers used to examine gene expression by RT-qPCR.

Accumulated SA and SA analytes were extracted and analyzed following the protocol and conditions described in [23]. In brief, SA metabolites were extracted using 20 mg of ground leaves and each sample was mixed with 0.25 mL of ice-cold methanol/water solution (20/80, v/v), including three internal standards (0.01 μg/mL salicylic acid-d_6_, for salicylic acid; 0.4 μg/mL 2,5-dihydroxybenzoic acid-d_3_, and 2,3-dihydroxybenzoic acid; and 0.8 μg/mL methyl salicylate-d_4_ for methyl salicylate). Samples were processed by ultra-sonic assisted extraction for 30 min in an ice block-filled bathtub. After incubation, samples were centrifuged at 30,000 g at 4°C during 10 min, the supernatant was recovered and filtered with a 0.22 μm membrane filter, and 10 μL was injected into LC–MS/MS system. LC–MS/MS analyses were carried out with an Ultimate 3000 LC system coupled to a TSQ Quantiva triple quadrupole mass spectrometer (Thermo Fisher Scientific, San Jose, CA, USA). The analytes (salicylic acid 2-O-β-D-glucoside, 2,3-dihydroxybenzoic acid, salicylic acid, and methyl salicylate) were chromatographed on a Thermo Fisher scientific Acclaim C30 column (150 mm × 2.1 mm, 3.0 μm particle size) at a column temperature of 30°C using a gradient elution with 0.1% formic acid in water (eluent A) and 0.1% formic acid in acetonitrile (eluent B). The gradient was as follows: 0-10 min 20-95% B and 10-15 min 95% B. The column was re-equilibrated using the initial mobile phase before next run. The flow rate was set at 0.2 mL/min. The mass spectrometer was operated in both positive and negative electrospray ionization (ESI+ and ESI−) with selected reaction monitoring (SRM) mode. The analytes were assigned by comparing SRM transitions and retention times with authentic standards. Xcalibur software (Ver. 3.0) was employed for data processing and instrument control.

#### Statistical analyses

To compare the amount of *C*Las DNA in mature leaves between inoculation treatments, and to determine the differences in gene expression and accumulation of SA and its metabolites, repeated measures analysis of variance (ANOVA) with Tukey’s test post hoc analyses were performed. All statistical analyses were run in RStudio environment [68].

## Acknowledgements

We are grateful to Dr. Kirsten Pelz-Stelinski for providing the equipment to perform molecular analyses. We thank Kristin Racine, Kara Clark, Angelique Hoyte and Rosa Johnson for their assistance in insect-plant care and data collection.

## Supporting information

**S1 Fig.**
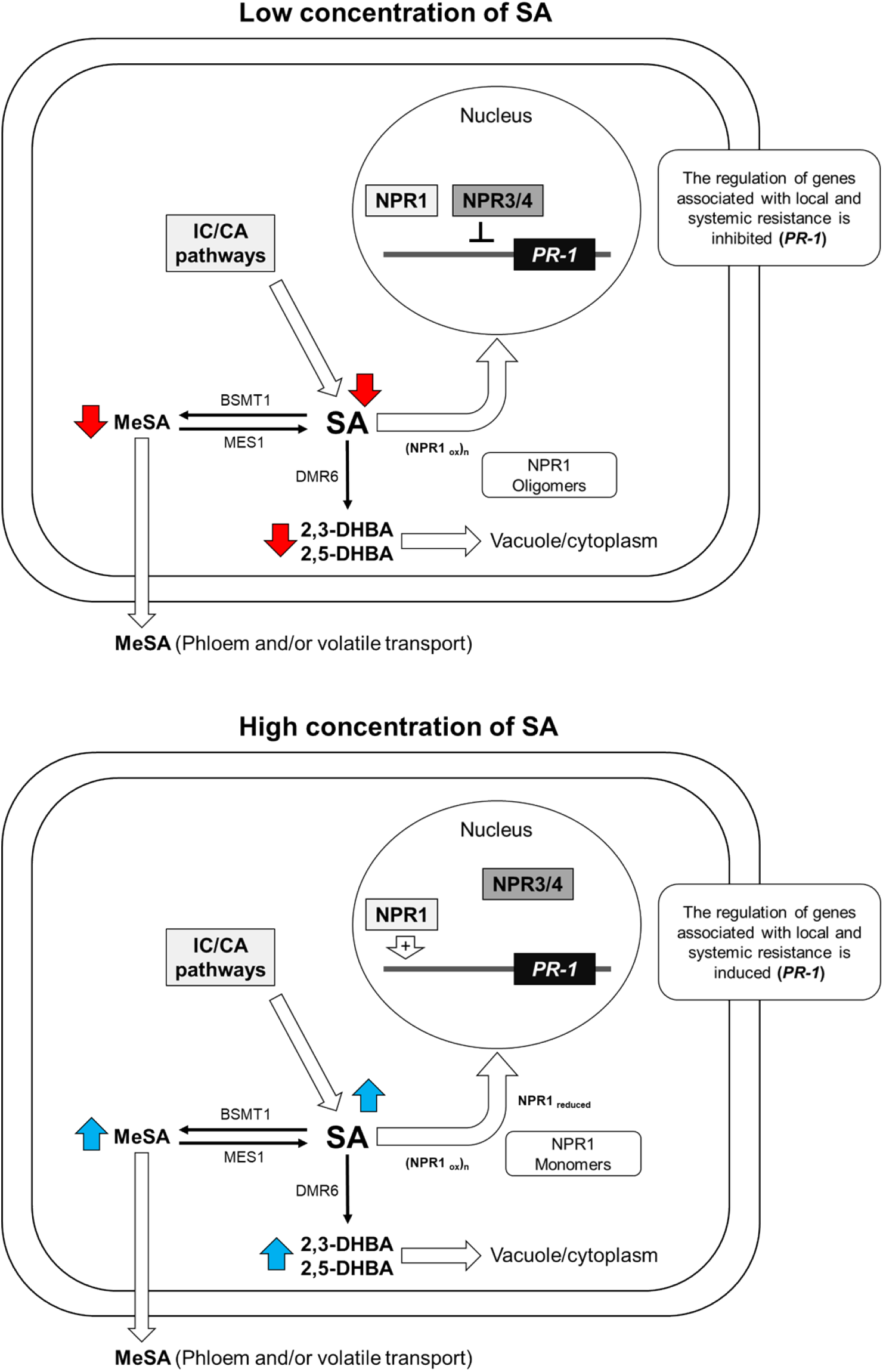
Proposed models of SA-dependent immune responses in *C. sinensis*.

